# Structural propensity database of proteins

**DOI:** 10.1101/144840

**Authors:** Kamil Tamiola, Matthew M Heberling, Jan Domanski

## Abstract

An overwhelming amount of experimental evidence suggests that elucidations of protein function, interactions, and pathology are incomplete without inclusion of intrinsic protein disorder and structural dynamics. Thus, to expand our understanding of intrinsic protein disorder, we have created a database of secondary structure (SS) propensities for proteins (dSPP) as a reference resource for experimental research and computational biophysics. The dSPP comprises SS propensities of 7,094 unrelated proteins, as gauged from NMR chemical shift measurements in solution and solid state. Here, we explain the concept of SS propensity and analyze dSPP entries of therapeutic relevance, *α*-synuclein, MOAG-4, and the ZIKA NS2B-NS3 complex to show: (1) how propensity mapping generates novel structural insights into intrinsically disordered regions of pathologically relevant proteins, (2) how computational biophysics tools can benefit from propensity mapping, and (3) how the residual disorder estimation based on NMR chemical shifts compares with sequence-based disorder predictors. This work demonstrates the benefit of propensity estimation as a method that reports both on protein structure, lability, and disorder.

## Introduction

Protein sequence is believed to hold the key to a perplexing mystery in the life sciences-the protein folding problem (***Dobson, 2003***). Therefore, immense efforts have been devoted to unraveling the sequence-structure relationship in polypeptides (***Baker, 2000; Bowie, 2005; Huang et al., 2016***). Although the fundamental forces of protein folding are known (***Dobson and Karplus, 1999; Karplus and Weaver, 1976; Karplus and Kuriyan, 2005***), complexity has hampered development of accurate folding prediction methods (***Moult et al., 2014***). Computational analysis of public protein databases, especially the Protein Data Bank (PDB) (***Varadi et al., 2015***), has played an integral role in shaping our fundamental understanding of protein structure, and for the advancement of protein design and structure prediction methodology (***Mackenzie and Grigoryan, 2017***). With accumulating structural data, it has become possible to mine for more complete and complex observations, which capture recurring structural features of proteins along with their sequence preferences. However, as explained in the seminal works of Dyson, Wright (***Dyson and Wright, 2004***) and Dobson (***Dobson, 2003***), naturally occurring protein disorder severely limits three-dimensional structure determination using X-ray crystallography, which renders only a rudimentary knowledge of the conformational state of disordered protein regions. Consequently, databases of protein structures are devoid of representative experimental data for intrinsically disordered proteins (IDPs) (***He et al., 2009***).

IDPs are abundant and control a vast array of biologically important processes, effectively complementing the functional spectrum of ordered proteins (***Dobson, 2003; Xie et al., 2007; Vucetic et al., 2003***). The prevalence of functional protein disorder (***Wright and Dyson, 2014***) demands reevaluation of the classical paradigm that a given protein sequence corresponds to a defined structure and function. Importantly, literature suggests that the biophysical features of IDPs and their protein interactions vary tremendously, and that there may be no common mechanism that can explain the different binding modes observed experimentally. Disordered proteins appear to make combined use of features such as pre-formed structure and flexibility, depending on the individual system and the functional context ***Mollica et al. (2016)***.

With no simple physical model that relates residual disorder to protein sequence and function, machine learning (ML) offers hope for unraveling the biophysical features of disordered protein regions (***Varadi et al., 2015; Dosztányi and Tompa, 2017; Hanson et al., 2016***). However, the quality and annotation level of input data will dictate the broad applicability of ML-based prediction tools, which are currently hindered through the incomplete implementation of two fundamentals factors: (1) sensitivity to intrinsic protein disorder at the residue level and (2) experimental conditions. For the latter, fundamental laws of equilibrium thermodynamics prove that experimental conditions influence protein structure and dynamics (***Finkelstein and Badretdinov AYa, 1997***). Numerous protein models in PDB database contain ambiguous disordered regions, where more than one structure of the same protein sequence "disagrees" in terms of the presence or absence of missing residues. A thorough survey of intrinsically disordered protein regions (IDPRs) suggests that ambiguity is a natural result for many proteins crystallized under different conditions. It is likely that structural ambiguity arises because many of intrinsically disordered protein regions were conditionally or partially disordered (***DeForte and Uversky, 2016***). Although specialized databases of intrinsically disordered proteins (IDPs) exist (***Varadi and Tompa, 2015; Piovesan et al., 2017; Yu et al., 2017***), they contain approximately only 900 fully annotated proteins with binary assignment of structural disorder, as gauged from coarse experimental techniques (***Dosztányi and Tompa, 2017***). However, there is a constant need for comprehensive and residue-specific datasets, which would enhance our understanding of intrinsic protein disorder and propel the development of better predictive methods.

Among the available experimental techniques, NMR spectroscopy has proven to be unique in its capacity to study disordered and folded proteins with atomic detail, both in solid and solution states***Felli and Pierattelli (2015)***. NMR chemical shifts are perfectly suited to help answer such questions, since they reflect the conformational preferences of polypeptide chains with atomic resolution***Wishart and Case (2001)*** and display exquisite sensitivity to local dynamics***Berjanskii and Wishart (2007); Wishart et al. (1992); Marsh et al. (2006)***. Furthermore, chemical shifts are easy to measure experimentally and can be efficiently assigned to individual atoms in the protein molecule.

To advance computational methodologies that are sensitive to experimental conditions and intrinsic disorder at the residue level, we have constructed a database of structural propensities of proteins (dSPP). Our repository is derived from a subset of 7094 NMR resonance assignments of unrelated proteins in solution and solid state near physiological conditions. The transpiring chemical shifts are perfectly suited to address the above issues in computational predictions, since they reflect conformational preferences of polypeptide chains (***Wishart and Case, 2001***) with high sensitivity to local dynamics (***Berjanskii and Wishart, 2007; Wishart et al., 1992; Marsh et al., 2006***). The dSPP database makes use of an enhanced version of the structural propensity method, which has been specifically developed for IDPs (***Tamiola and Mulder, 2012***), thus providing optimal sensitivity to residual disorder for folded and unstructured polypeptides.

To demonstrate the value in dSPP, we first compare the structural propensity mappings from dSPP with corresponding 3D structures of therapeutically-relevant database entries, *α*-synuclein (*α*S), its aggregation modifying protein (MOAG-4), and NS2B-N3S protease complex from the Zika virus (ZIKV). Subsequently, we explain the relative benefit of structural propensity mapping in machine learning methodology. Lastly, empirically derived disorder scores from ZIKV are compared with theoretical disorder predictions from six state-of-the-art tools (***Hanson et al., 2016; Ishida and Kinoshita, 2007; Linding et al., 2003; Ward et al., 2004***). Our work concludes with discussing how structural propensity data can propel development of structure and disorder prediction tools with higher accuracy and computational efficiency.

## Results

### NMR Assignment Data

A subset of 11286 protein resonance assignment entries was retrieved from the Biological Magnetic Resonance Data Bank (***Ulrich et al., 2008***). Within the downloaded records, 3286 contained more than one resonance assignment. Upon stringent filtering, 5860 assignment records were rejected because of a suboptimal length, 2426 were omitted due to the lack of ^13^*C^α^* and ^13^*C^β^*, and 335 entries were removed that contain abnormal backbone resonance assignments due to the use of paramagnetic agents, non-protein compounds, and assignment errors. The final dataset consisted of 7094 protein resonance assignments. The average level of resonance assignment completeness (Supplementary Table 2) for ^1^*H^N^*, ^1^*H^α^*, ^13^*C^α^*, ^13^*C^β^*, ^13^*C^O^*, and ^15^*N* was found to be 79%.

### Experimental Conditions

Experimental parameters having profound biophysical effects on protein structure were assessed for the dSPP entries: temperature, pH, and ionic strength. Figure 1 plots the distributions of experimental conditions in dSPP. Panel 1a shows that temperature centers around 296K ± 10 K(min., 283 K; max., 323 K). Panel 1bshowsthata majority of database entries were studied in a physiological pH range with a mean pH of 6.9 ± 0.7. Furthermore, panel 1c shows a dominant contribution of low-salt resonance assignments that center around an ionic strength of approx. 82 mM ± 87 mM. Despite efforts towards standardization of data referencing (***Wishart et al., 1995***), NMR spectra are known to contain systematic referencing errors (***Zhang et al., 2003***). Panel d reflects this notion by reporting a mean offset correction of 0.36 ppm and standard deviation of 0.02 ppm, which follows a unimodal distribution.

**Figure 1.**
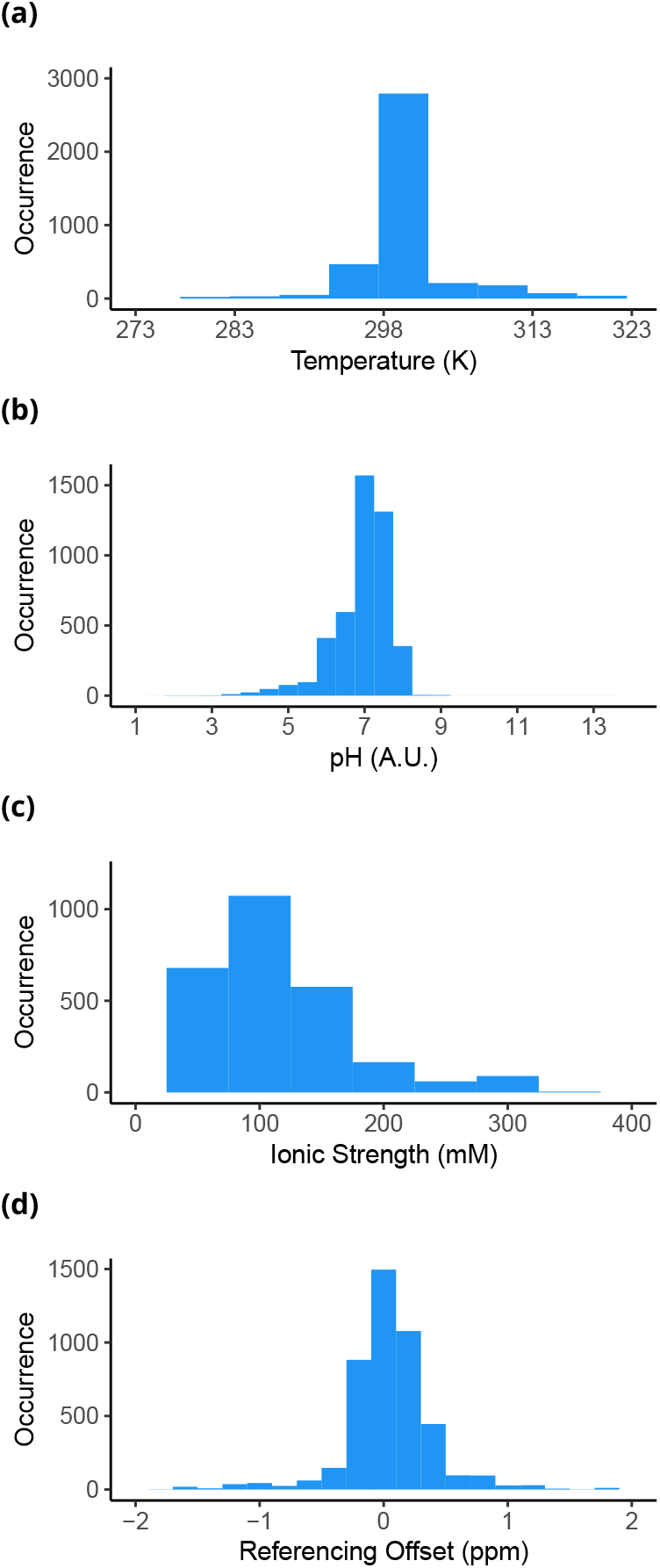
Statistical summary of experimental conditions in dSPP. Frequency distribution plots for (a) temperature, (b) pH, (c) ionic strength, and (d) resonance assignment referencing offset.

### Protein Sequence Statistics

As demonstrated in Figure 2a, dSPP contains 7094 protein sequences with the mean length of 119 ± 61 residues distributed in a unimodal fashion. Upon sequence homology and residue conservation analyses (Methods), the mean sequence conservation among aligned dSPP members is 0.11. Based on Figure 2b, it is apparent that amino acids are not distributed uniformly. Other protein sequence databases generate the same trend, which has been extensively studied from the perspective of sequence conservation analysis (***Valdar, 2002***). Dominant residues in dSPP are leucine, alanine, glutamate, and glycine; whereas cysteine and tryptophan are the least abundant.

**Figure 2.**
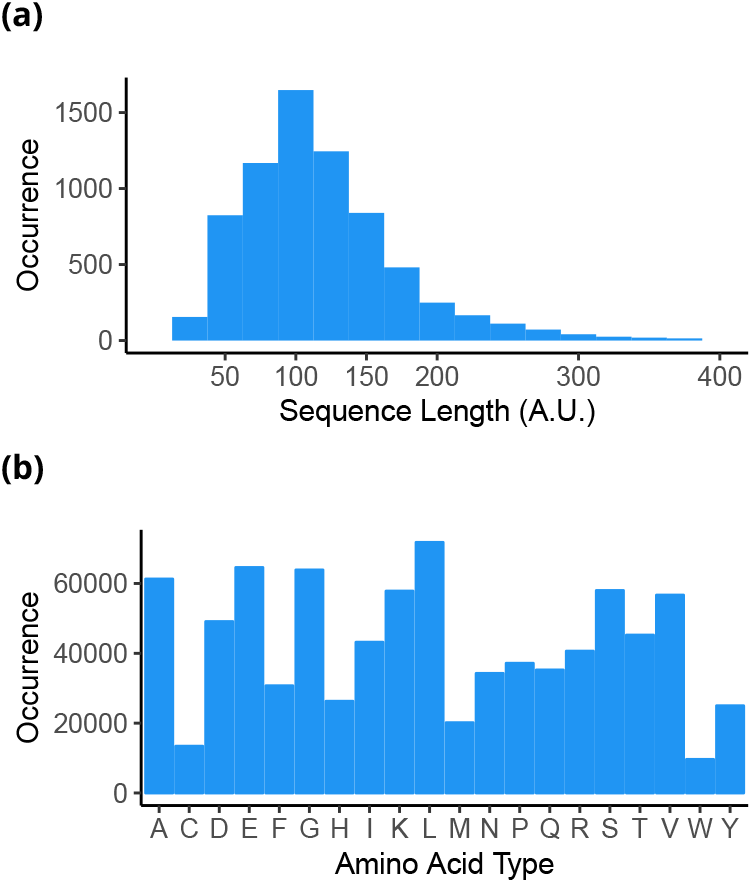
Statistical summary of protein sequences in dSPP. Frequency distribution plots for (a) protein sequence length and (b) amino acid composition.

### Structural Propensity Statistics

Structural propensity is derived from the differences between experimental backbone chemical shifts and empirically predicted shielding constants observed in IDPs of similar composition (***Tamiola et al., 2010***). Therefore, structural propensity (Ψ) is a measure of the departure of an individual polypeptide residue from canonical SSs towards a ‘random-coil’ state. Calculation of the residue-specific structural propensity score is described in the Methods section. We assume the ‘random-coil’ state can be modeled after the ensemble behavior and characteristics of IDPs in solution (***Tamiola and Mulder, 2012***). The structural propensity adopts real number values. A residual score of −1.0 indicates a fully formed *β*-sheet, whereas a propensity of 1.0 suggests that 100% of ensemble members at a given polypeptide position adopt an α-helical conformation. Importantly, a near-zero score (0.0) indicates residual behavior and conformational sampling observed in IDPs. Fractional propensity scores are quantitative indicators of structural lability. Therefore, a score of −0.5 or 0.5 signifies that 50% of ensemble members sample conformations that are neither *β*-sheet nor α-helix, respectively.

The profile of structural propensities in Figure 3a shows a dominance by disordered and near-helical segments in dSPP entries. The low abundance of near-*β* residual propensities is directly related to a sparse representation of all-*β* proteins in the BMRB database. Figure 3b provides fundamental evidence for conformational preferences of individual polypeptide residues in solution near physiological conditions. The skewed propensities towards disordered segments by glycine, proline, serine, and threonine is known (***Dwyer, 2006; Kumari et al., 2015***). Interestingly, a difference in structural preference is observed for aspartate and glutamate. Although both residues were reportedly abundant in disordered proteins (***Dyson and Wright, 2005; Uversky, 2016, 2014***), our analysis reveals a clear preference of aspartate (smaller side-chain) to populate disordered states, whereas glutamate displays a preference to populate more compact, a-helical conformations.

**Figure 3.**
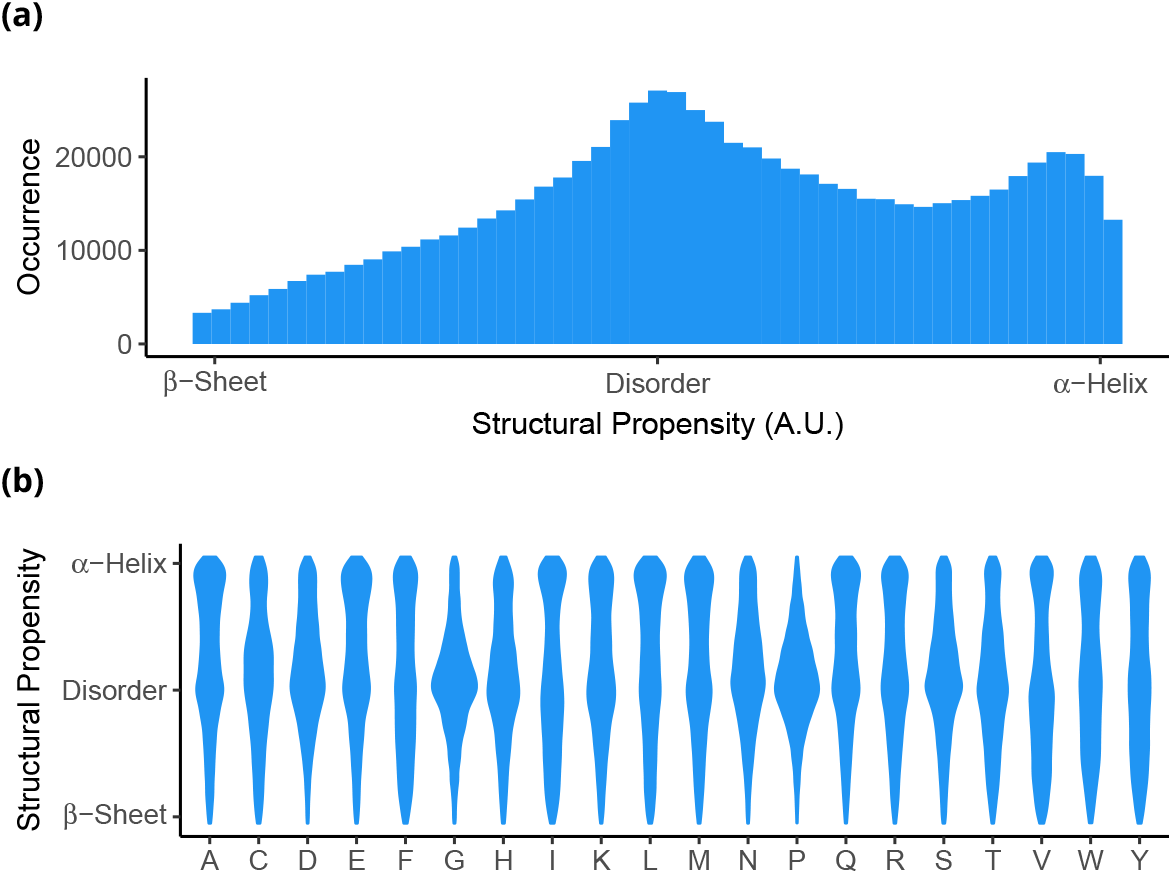
Statistical summary of structural propensity distribution in dSPP. Frequency distribution plots for (a) collective structural propensity and (b) normalized, residue-specific distribution of structural propensities. The widths of individual plots are directly proportional to the distribution density at the given structural propensity value. .

### Examples of Structural Propensity Mapping

Structural propensities are directly mapped to atomistic structures derived from either NMR restraints or X-ray crystallography, where both have supplementary dynamics studies and extensive biophysical characterization. The models αS, MOAG-4, and ZIKV NS2B-NS3 complex are prime targets for drug discovery. Subsequently, we demonstrate the practical advantage of structural propensity over canonical structure-based classification methods and demonstrate how six seminal disorder predictors score against experimentally derived structural propensities for ZIKV NS2B-NS3 complex.

#### Intrinsically disordered a-synuclein

*α*-synuclein (*α*S) is a 140-residue IDP with high net charge and low hydrophobic content that has been implicated in a vast array of highly debilitating neurodegenerative conditions; most notably the Parkinson’s disease (PD) (***McCormack and Di Monte, 2009; Luk et al., 2012***). *α*S is believed to be involved in the regulation of the homeostasis of synaptic vesicles during neurotransmitter release (***Cooper et al., 2006***) and it has been suggested to play a crucial role in the interactions with vesicular membranes in both physiological and pathological contexts (***Cooper et al., 2006***). Since a hallmark of PD is the formation of abnormal intracellular protein aggregates of αS, referred to as Lewy bodies (***Cooper et al., 2006***), αS and its biophysical characterization have become the focal point of research on IDPs and neuropathology. Figure 4a shows the structural ensemble model of αS derived from experimental restraints; NMR chemical shifts and paramagnetic relaxation enhancement of NMR measurements (***Fusco et al., 2016***). As evidenced by supporting experimental data, under near-physiological conditions αS exists as a highly labile entity that inter-converts between a multitude of conformations. This notion is reflected in ensemble averaged secondary structure (SS) fraction depicted in Figure 4b, which demonstrates a lack of persistent canonical SS preference in αS ensemble. The structural propensity for αS, adopted from dSPP entry dSPP19351_0, is given in Figure 4c. Our structural propensity analysis displays good qualitative agreement with the ensemble model of αS. However, upon closer inspection, propensity scores for αS reveal the existence of three structural domains (***Luk et al., 2012***): 1) an N-terminal domain (residues 1-60) that supports regions of transient α-helical propensity; 2) a central hydrophobic region, known for historical reasons as the “non-Amyloid *β* component” (NAC) (residues 61-95) that is itself highly amyloidogenic and forms the core of the amyloid fibrils (***Vilar et al., 2008***); and 3) a C-terminal, acidic and proline-rich segment with residual tendencies to adopt extended structures (residues 96-140).

**Figure 4.**
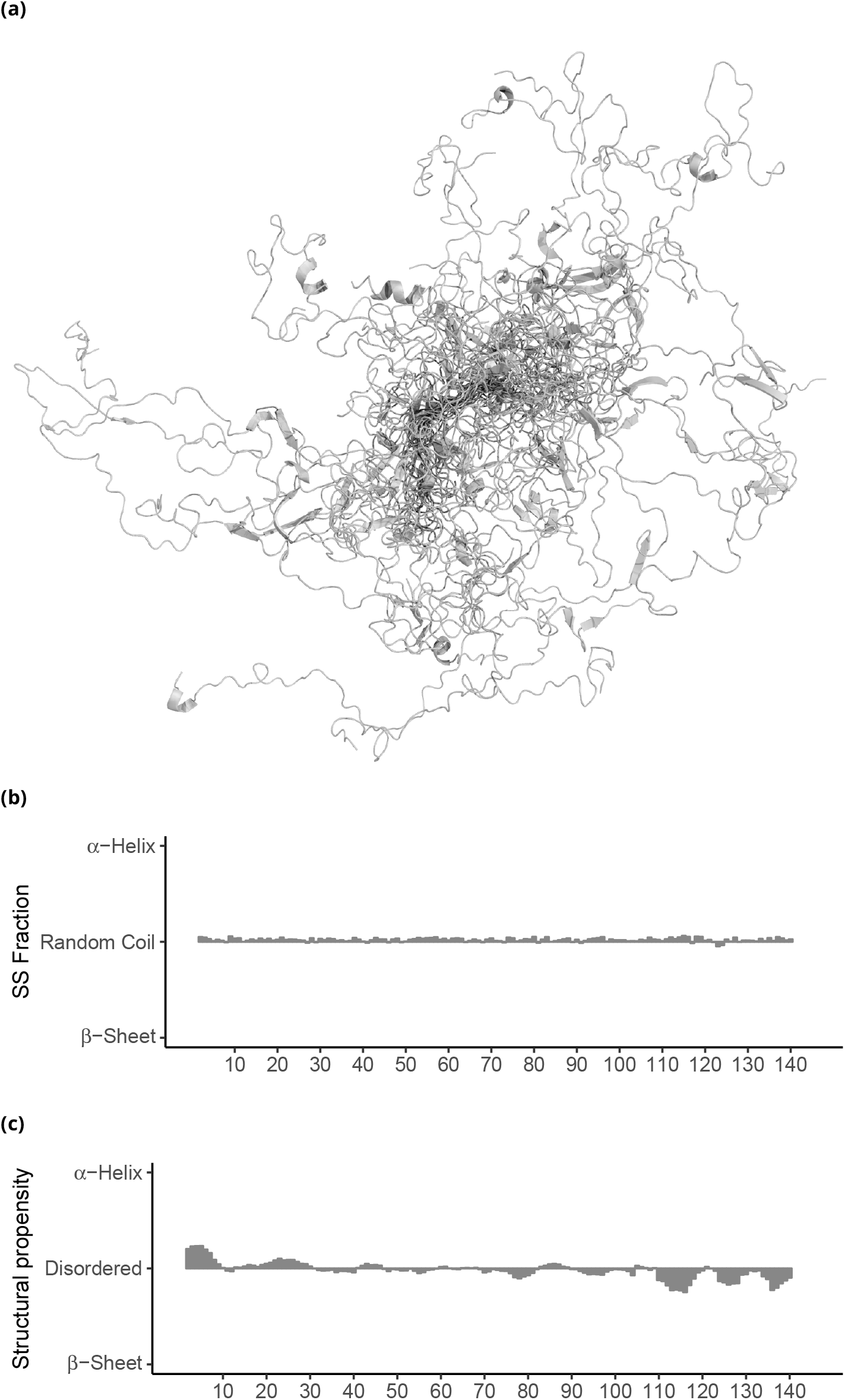
Structural ensemble of human **α**-synuclein. (a) Superposition (using 10-30) of fifty low-energy conformers from ensemble of **α**S (PED9AAC). (b) Residue-specific secondary structure (SS) fraction computed from models present in (PED9AAC) ensemble. (c) Structural propensity mapping for **α**S from dSPP database (dSPP19351_0).

#### Partially disordered Modifier of Aggregation 4 (MOAG-4)

Although biophysical features and ensemble properties of *α*S have been elucidated, the regulatory forces behind *α*S aggregation in the cellular environment remain elusive. A study of *α*S aggregation behavior in *C. elegans* models led to the discovery of a regulator of protein aggregation; ‘modifier of aggregation 4’ (MOAG-4) (***van Ham et al., 2010***). It has been shown that inactivation of MOAG-4 leads to suppression of protein aggregation and associated toxicity in *C. elegans* models for *α*S (***van Ham et al., 2010***). Importantly, the human ortholog of MOAG-4, EDRK-rich factor (SERF) 1A, accelerates the aggregation of a broad range of amyloidogenic proteins in vitro in the initial stage of the process (***Falsone et al., 2012***). In a recent and extensive study, ***Yoshimura et al. (2017)*** investigated the kinetic and structural effects of MOAG-4 on the aggregation of *α*S. Figure 5a shows the structural ensemble of MOAG-4 ***Yoshimura et al. (2017)***, calculated from NMR backbone chemical shifts (***Camilloni et al., 2013***) and cross-validated by prediction of experimental NOEs (***Yoshimura et al., 2017***). As evidenced by ensemble-average fractional SS analysis in Figure 5b, MOAG-4 is a partially disordered polypeptide with two distinct regions of *α*-helical propensity: lowly-populated, transient *α*-helix comprised of residues 7-20, and well-defined helical segment made of residues 39-70. The structural propensity mapping from NMR resonance assignments for MOAG-4 is available in dSPP as dSPP18841_0 and shown on Figure 5c. Computed structural propensity displays an excellent quantitative agreement with fractional SS, clearly demonstrating transient character of the 7-20 helical segment relative to the defined *α*-helix of 39-70 fragment.

**Figure 5.**
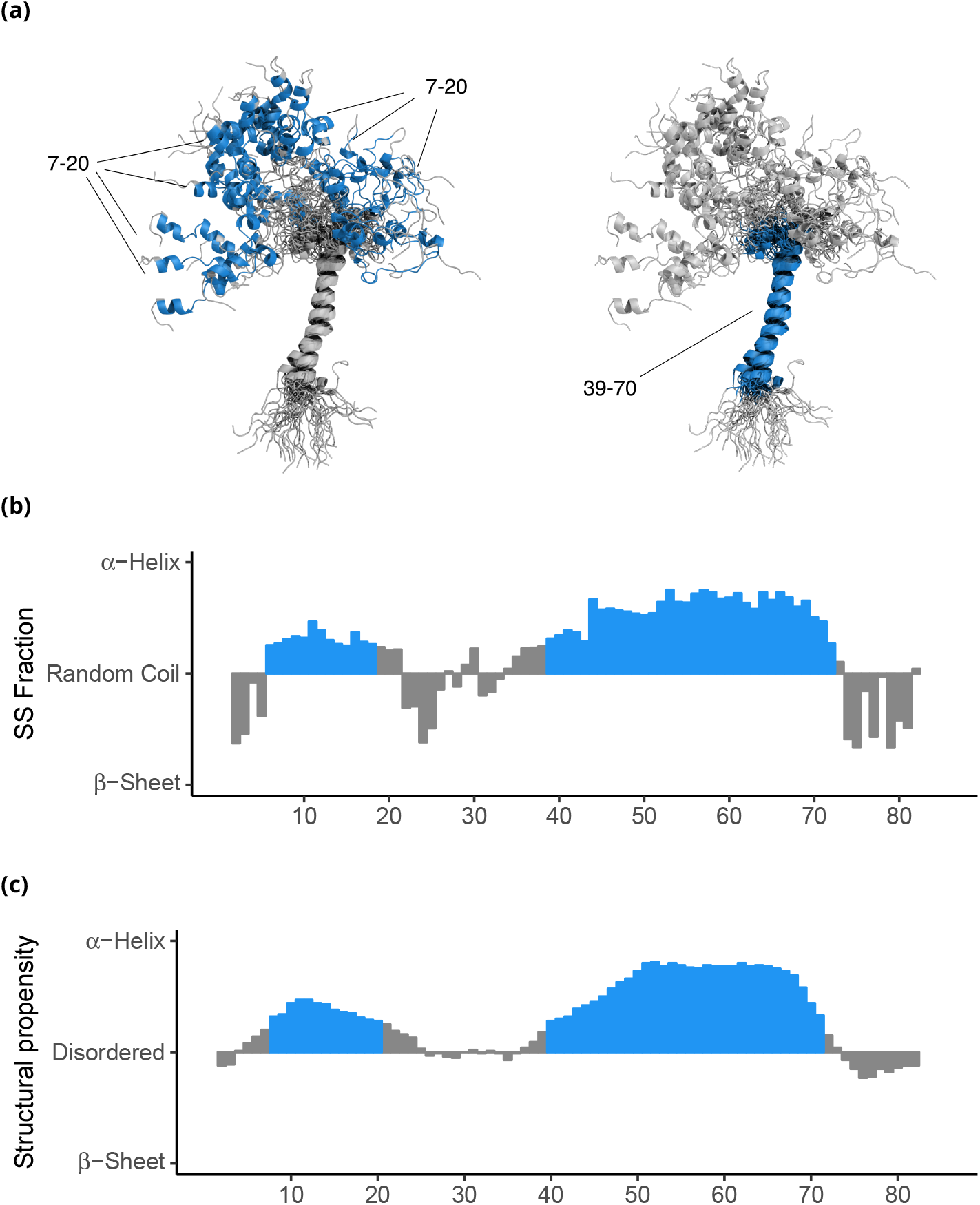
Structural ensemble of MOAG-4. (a) Fifty low energy conformers from ensemble of MOAG-4. (b) Ensemble-averaged secondary structure (SS) fraction computed from models present in PED5AAA ensemble. (c) Structural propensity from dSPP database (dSPP18841_0). The segments of MOAG-4 with SS fraction and propensity higher than 0.25 are marked in blue.

#### NS2B-NS3 protease from Zika virus

The Zika virus (ZIKV) is a highly contagious representative of pathogenic flaviviruses and is linked to fetal microcephaly and neurological complications in adults, such as Guillain-Barré syndrome, acute myelitis, and meningoencephalitis (***Petersen et al., 2016***). The flavivirus NS2B-NS3 protease is a main target for antiviral therapeutics due to its role in ZIKV replication (Luo et al., 2015). Recently, NMR resonance assignments and crystal structures of the NS2B peptide cofactor complexed with NS3 protease from ZIKV (PDB: 5GJ4)were reported (***Zhang et al., 2016***). Structural studies of the NS2B-NS3 complex have been complemented with an analysis of ^15^*N T*_1_, *T*_2_ NMR relaxation times and hetNOE experiments in solution (***Zhang et al., 2016***). The structural propensity and experimental conditions for the NS2B peptide and NS3 protease are available in dSPP under accession numbers dSPP26928_0 and dSPP26928_1, respectively. The structural propensity analyses of the NS2B-NS3 complex in Figures 6a and 6b show that both NS2B and NS3 display a heterogeneous distribution of structural disorder. Both N- and C-termini of NS3 protease resemble structural disorder found in IDPs, which are in an excellent agreement with the reported ^15^*N T*_1_, *T*_2_, and hetNOE NMR experiments. It has been shown that residues 1-17 and 170-177 of the respective N- and C-termini in the NS3 protease are highly dynamic in solution, as evidenced by low *T*_1_ and hetNOE values (< 0.6) (***Petersen et al., 2016***). As a result of extensive dynamics, both termini are missing in the X-ray structure. Conversely, although NS2B induces a closed complex with NS3 protease in solution, our structural propensity analysis indicates that the C-terminal of NS2B and residues around the P2 catalytic pocket in NS3 exhibit a large degree of disorder, which is contrary to the X-ray structure. Additionally, residues 80-95 of NS2B, which form a *β*-hairpin in the crystal structure, were found to be highly dynamic in solution, as evidenced by severe NMR spectral broadening and negative hetNOEs (***Zhang et al., 2016***). Thus, the NS2B-NS3 complex displays completely different dynamics near physiological conditions (ionic strength: 170 mM; pH: 7.3; pressure: 1 atm; temperature: 310 K) in solution-state NMR experiments compared to the conditions in X-ray crystallography (0.2M Sodium Malonate pH 4.0; 20% PEG 3350; flash-frozen in liquid *N*_2_).

**Figure 6.**
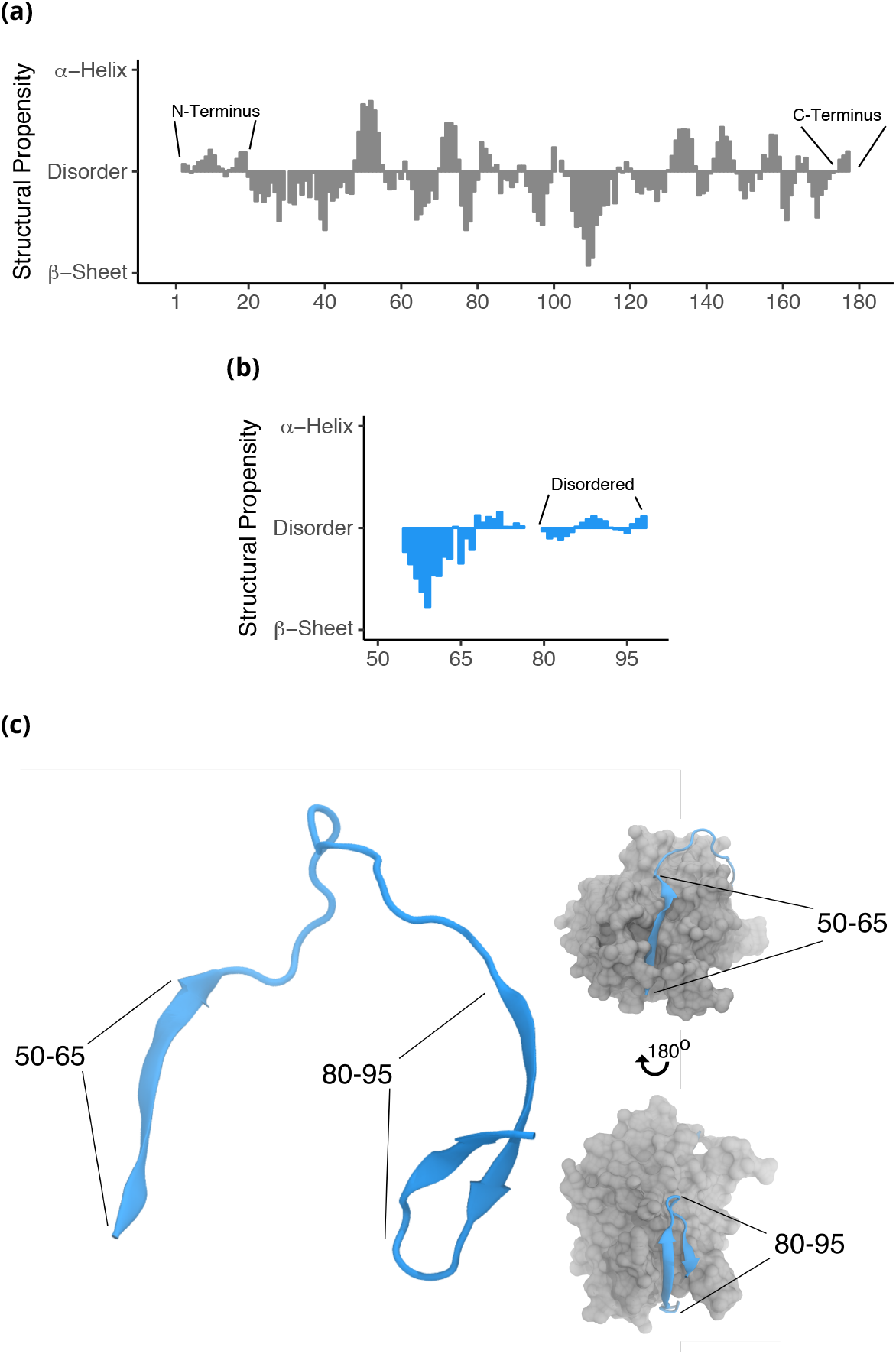
Structural propensity analysis for ZIKV NS2B-NS3 protease complex. Structural propensity plots for (a) NS3 protease and (b) NS2B peptide. The disordered segments of NS2B and NS3 are marked accordingly. (c) Comparative assessment of structural propensity and X-ray crystallography model (PDB: 5GJ4) for ZIKV NS2B-NS3 complex. The structure of NS2B polypeptide is given in blue, whereas NS3 is depicted in gray. Left-hand panel depicts cartoon representation of NS2B with 50-65 and 80-95 beta-hairpin segment, as gauged from X-ray model. The right-hand side panel demonstrates close-complex of NS2B-NS3.

#### Structural propensity as an input for machine learning

The relative benefit of machine learning as a computational mean for protein structure and disorder prediction is its ability to accept arbitrary input and output data types (***Wang et al., 2016***). ML methods aimed specifically at structural disorder prediction are predominantly trained on datasets of binary-encoded, multi-class SS types. Protein disorder is inferred from missing structural coordinates, sequence conservation, similarity to known disordered proteins, and 3D contact maps (***Wang et al., 2017; Jones, 1999; Ishida and Kinoshita, 2007***). The assignment of *c* canonical classes of SSs for a peptide of *N* amino acids translates into a ‘one-hot’ encoded *c* × *N* tensor *X_c_* which is then used as input for ML, as given by Equation 1.

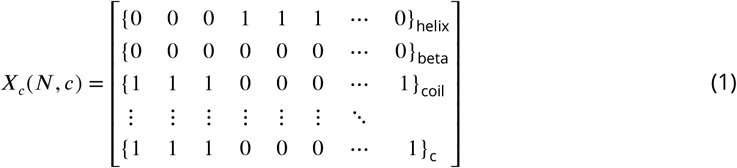

The structural propensity approach offers a computationally effective alternative to binary-type SS class assignments. The structural features of a polypeptide chain from Equation 1, can be encoded using a 1 × *N* tensor *X*_Ψ_ shown in Equation 2.

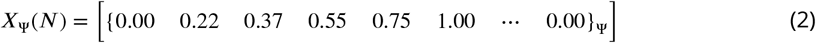

The computational gain due to replacement of binary-class assignment by structural propensity is directly proportional to the number of structural classes *c*. Thus, structural propensity representation of a triple-class tensor reduces parameter search space by three-fold, which translates into shorter training time and better convergence (***Qian and Sejnowski, 1988***).

### Comparison of disorder predictions and absolute experimental propensity scores

We have predicted structural disorder probabilities for the ZIKV peptides using SPOT (***Hanson et al., 2016***), PrDOS (***Ishida and Kinoshita, 2007***), variants of DisEMBL; Loops, Hot Loops, Remark465 (***Linding et al., 2003***) and Disopred 3 (***Ward et al., 2004***). The disorder probability predictions were compared against the absolute structural propensity score Ψ_*abs*_, which reports on a normalized probability that residues within the polypeptide chain sample ‘random-coil’ conformations. Ψ_*abs*_ is computed from Ψ_*abs*_ = 1 − |Ψ|, and shown in Figure 7 in blue. Major discrepancies in the magnitude of predicted disorder probabilities are apparent for all theoretical methods. Smoothly interpolated disorder predictions by SPOT match only the fragments of N- and C-terminally disordered segments of NS2B-NS3. Compared to the experimentally derived Ψ_*abs*_ scores, SPOT displays the smallest sensitivity to structural detail among all computational methods. SPOT cannot discern stable SSs from confirmed domains of structural disorder in all tested polypeptides. Relative to SPOT, PrDOS produces more dispersed probabilities, yet still at great variance with Ψ_*abs*_. The systematic underestimation of structural disorder using DisEMBL Hot Loops method is also apparent, which is accentuated in Figure 7 where the method assigns near-zero disorder probability for the entire NS3 protein. This finding not only contrasts with the Ψ_*abs*_ score and NMR data, but disagrees profoundly with the trends reported by remaining predictors. Remark 465 and Loops display limited sensitivity to residual structure. As shown in Figure 7, DisEMBL Loops consistently produces probability distribution, which qualitatively resembles experimental Ψ_*abs*_, yet differs significantly in the magnitudes of predicted disorder probabilities.

**Figure 7.**
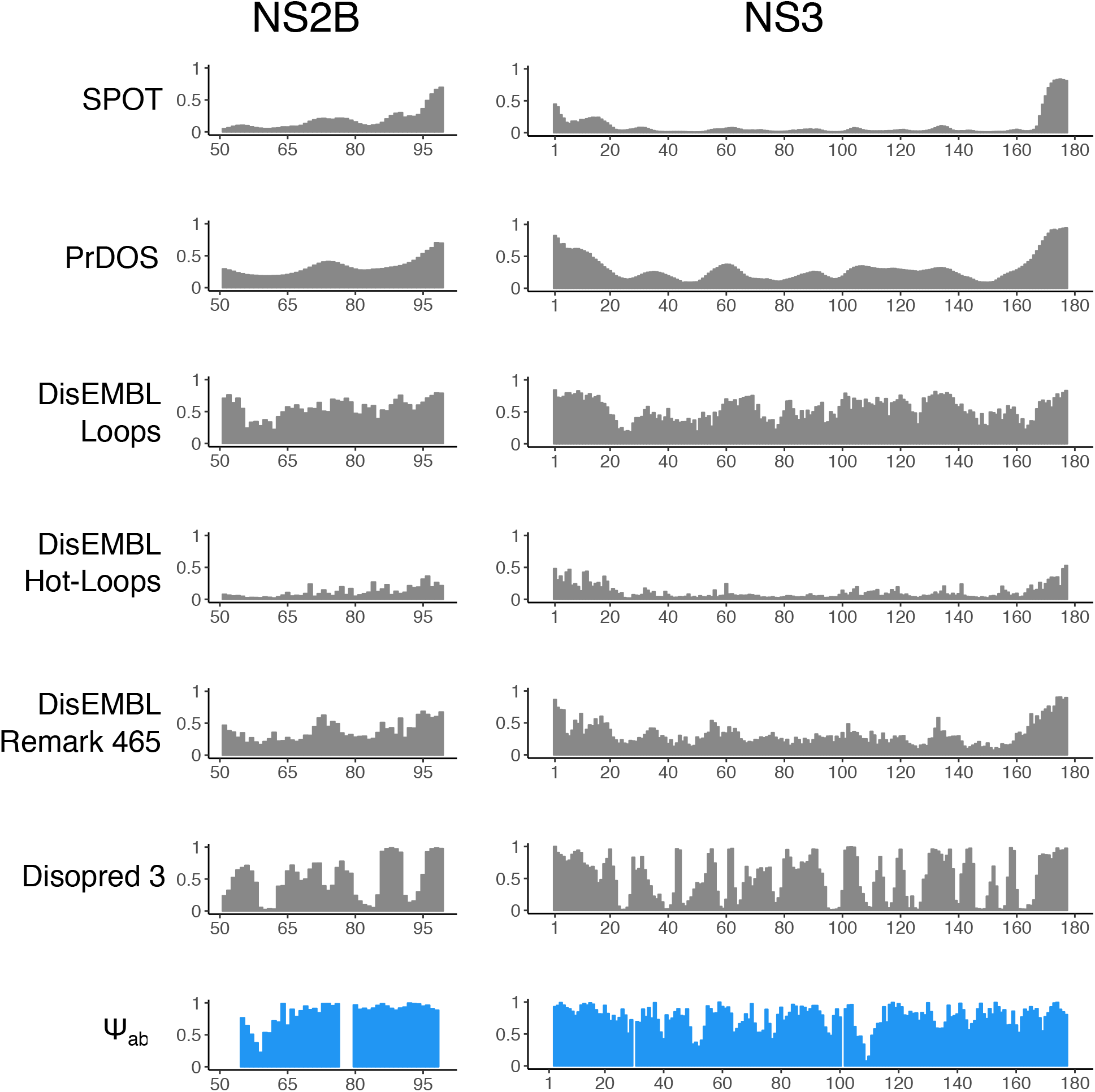
Prediction of probability of structural disorder using SPOT, PrDOS, DisEMBL Loops, Hot-Loops, Remark 465, Disopred 3 and absolute experimental structural propensity score Ψ_*abs*_, shown in blue.

Table 1 contains the outcome of non-parametric single-way ANOVA analyses of computed disorder probability distributions with respect to experimental absolute structural propensity score Ψ_*abs*_. The Disopred 3 method consistently produced the distribution of disorder scores with the smallest disparity between the experimental propensity. The method managed to properly assign structural disorder to N- and C-terminal domains of NS3, as well as properly identify *β*-extended 55-65 segment of NS2B. However, Disopred 3 systematically underestimated structural disorder in both NS2B and NS3 peptides. This notion was reflected by low value of χ^2^ and *p*. A general inspection of power divergence χ^2^ and distribution similarity probability p clearly suggest that all theoretical tools yield disorder probability distributions that are different from experimental absolute structural propensity scores for NS2B and NS3 proteins.

**Table 1.**
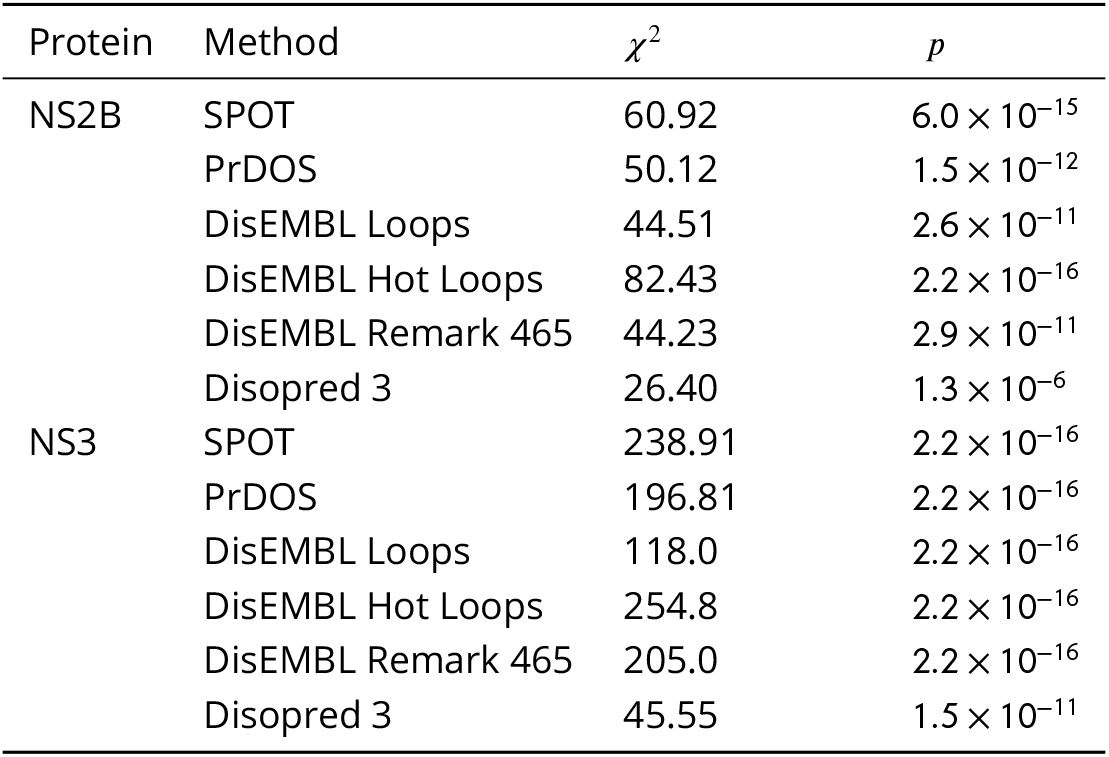
Non-parameteric single-way ANOVA (Kruskal-Wallis rank sum test) analyses of disorder prediction distributions for ZIKV (NS2B and NS3) proteins using SPOT, PrDOS, DisEMBL Loops, DisEMBL Hot Loops, DisEMBL Remark465 and Disopred 3 against absolute experimental propensity Ψ_*abs*_. χ^2^ denotes power divergence of predicted data with respect to experimental propensities, and *p* is the probability of distribution similarity. The ANOVA test was performed assuming one degree of freedom.

| Protein | Method | $\chi^2$ | $p$ |
| --- | --- | --- | --- |
| NS2B | SPOT | 60.92 | $6.0 \times 10^{-15}$ |
| | PrDOS | 50.12 | $1.5 \times 10^{-12}$ |
| | DisEMBL Loops | 44.51 | $2.6 \times 10^{-11}$ |
| | DisEMBL Hot Loops | 82.43 | $2.2 \times 10^{-16}$ |
| | DisEMBL Remark 465 | 44.23 | $2.9 \times 10^{-11}$ |
| | Disopred 3 | 26.40 | $1.3 \times 10^{-6}$ |
| NS3 | SPOT | 238.91 | $2.2 \times 10^{-16}$ |
| | PrDOS | 196.81 | $2.2 \times 10^{-16}$ |
| | DisEMBL Loops | 118.0 | $2.2 \times 10^{-16}$ |
| | DisEMBL Hot Loops | 254.8 | $2.2 \times 10^{-16}$ |
| | DisEMBL Remark 465 | 205.0 | $2.2 \times 10^{-16}$ |
| | Disopred 3 | 45.55 | $1.5 \times 10^{-11}$ |

## Discussion

X-ray crystallography and NMR spectroscopy have proven to be pivotal techniques in the determination of precise molecular models of proteins. However, polypeptides owe their complexity to an intricate combination of structure and dynamics (***Ishima and Torchia, 2000***). A dynamic view of protein structures can rationalize effects from experimental conditions or structural features and disorder that result in altered function and pathology. Understanding the link between protein structure and dynamics is particularly important in proteins with intrinsically disordered domains, which are prevalent in nature, have been implicated in numerous human pathologies, and are difficult to study by conventional structural biology methods (***Vucetic et al., 2003; Dobson, 2003; Uversky et al., 2008; Wright and Dyson, 2014; Uversky, 2014***). We have shown here how the inclusion of NMR-derived structural propensity can enhance biological understanding from static crystal structures, provide extensive structural characterization for IDPs involved in human pathologies, and assess the accuracy of computational methods that aim to discern stable SSs from disordered regions. This approach transpires from our dSPP public database, which should facilitate the inclusion of NMR-based SS descriptors in existing structure and disorder prediction methods and serve as a resource for structural interpretation of protein NMR chemical shifts.

The exquisite sensitivity of SS propensity to structural detail at the residue level is reflected in our assessment of intrinsically disordered α-synuclein (***Fusco et al., 2016***) and partially folded MOAG-4 (***Yoshimura et al., 2017***). We demonstrate that αS displays detectable propensity variation throughout its polypeptide chain, which corresponds to the presence of known micro-domains involved in αS function and aggregation behavior (***Luk et al., 2012***). Importantly, our propensity calculations agree well with the fractional SS estimation from structural ensemble models of αS computed using NMR chemical shifts and paramagnetic distance restraints (***Fusco et al., 2016***), reinforcing the notion that structural propensity reports on ensemble average properties of αS. Furthermore, the comparative assessment of structural features of MOAG-4 ensembles (***Yoshimura et al., 2017***) and SS propensity mapping clearly show that our method succeeds in structural characterization of polypeptides with heterogeneous distribution of structured and highly disordered domains. Additionally, ***Yoshimura et al. (2017)*** demonstrated that SS propensity could be used as a powerful agent to characterize structural effects of inter-molecular αS and MOAG-4 interactions, thus extending the application of SS propensity analysis to protein-protein complexes. Our SS propensity mapping of the ZIKV NS2B-NS3 protease complex shows that NS2B at near-physiological conditions exhibits a high level of structural disorder commonly found in intrinsically unstructured proteins (***Tamiola and Mulder, 2012; Tamiola et al., 2010***). Very good agreement between our analysis and hetNOEs NMR experiments (***Zhang et al., 2016***) suggests that the ordered *β*-hairpin motif in the NS2B X-ray model is a transient structure, which may obscure dynamics of the P2 interaction site at the NS2B-NS3 interface. As this interaction site has been designated as a potential drug design target (***Zhang et al., 2016***), NMR-derived structural propensities and relaxation techniques seem to be the most optimal tools to investigate changes in structure and dynamics of NS2B-NS3 upon drug binding under physiological conditions. The ultimate benefit of SS propensity as an experimental proxy for residual structure and disorder can be judged from our assessment of disorder predictors SPOT (***Hanson et al., 2016***), PrDOS (***Ishida and Kinoshita, 2007***), variants of DisEMBL; Loops, Hot Loops and Remark465 (***Linding et al., 2003***), and Disopred 3 (***Ward et al., 2004***). We show how absolute SS propensity can be used as a benchmark for residual disorder probability predictions, complementing existing approaches (***Moult et al., 2014***). Our analyses indicate that established disorder prediction methods suffer from insufficient sensitivity to disordered regions among folded domains. Our findings align very well with an extensive study by DeForte and Uversky (2016) who have shown that intrinsically disordered regions (IDRs) gauged from missing coordinates in PDB database may not be representative of intrinsic protein disorder. Thus, methods based of IDRs, including DisEMBL Loops, Hot Loops and Remark465 severely lack sensitivity to disorder behavior experimentally observed in IDPs. We postulate that our normalized SS propensity score could help to refine existing prediction methodologies and serve as an alternative to multi-dimensional, binary representation of protein structure classes for input in ML methods. This reduces numerical complexity and computational effort in development and training (***Dosztányi and Tompa, 2017***), and opens a possibility to include structural lability in predictive algorithms.

However, as a structure characterization technique based on chemical shift analyses, propensity mapping is subject to the theoretical and practical limitations of protein NMR spectroscopy (***Felli and Pierattelli, 2015***). The size of studied proteins and consequential overlap in spectral resonances, signal broadening, extensive chemical exchange, dynamics on different time scales (***Ishima and Torchia, 2000***), and the complex nature of electronic contributions to measured chemical shifts (***Wishart et al., 1992; Berjanskii and Wishart, 2006***) all limit the submission of NMR resonance assignments to public repositories. Furthermore, the current implementation our structural propensity model is fine-tuned to detect deviations from canonical *α*– or *β*– structures towards the disordered state only. It is expected that an expansion of the structural propensity concept to other SS classes (***Kabsch and Sander, 1983***) could further refine the computed SS propensity scores.

By transforming NMR resonance assignments of 7094 proteins at different experimental conditions into a database of structural propensities scores, we have created a resource that can enhance biological understanding of proteins with known NMR resonance assignments and propel the development of computational methods that aim to discern stable SSs from disordered regions. We hope that our structural propensity repository with a fully automated update cycle will benefit the machine learning community by providing a simple descriptor of SS class that is sensitive to structural disorder.

## Methods

### NMR Resonance Assignment Data

A subset of protein resonance assignment records with ^1^*H^N^*, ^1^*H^α^*, ^13^*C^α^*, ^13^*C^β^*, ^13^*C^O^*, and ^15^*N* nuclei was retrieved from Biological Magnetic Resonance Data Bank (***Ulrich et al., 2008***). Entries with sequence length of less than 30 amino acid residues and containing non-standard amino acids, nucleic acids, and paramagnetic agents were excluded from further analysis. The final screening for resonance assignment data was performed by selecting the resonance assignments which contained at least ^13^*C^α^* and ^13^*C^β^* nuclei, simultaneously with the absolute referencing offset smaller than |Δ_*r*_| < 2.0 ppm.

### Secondary Chemical Shift Calculations

The sequence-dependent deviations of experimental resonance assignments from the ‘random coil’ chemical shifts, known as secondary chemical shifts, were calculated using the ncIDP chemical shift library (***Tamiola et al., 2010***). In our procedure, the ‘random-coil’ chemical shift for a nucleus *n* ∈ { ^1^*H^N^*, ^1^*H^α^*, ^13^*C^α^*, ^13^*C^β^*, ^13^*C^O^*, ^15^*N* } of amino acid residue a, within a tripeptide *x* − *a* − *y*, is expressed as,

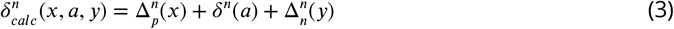

where *δ^n^*(*a*) is the ‘random-coil’ chemical shift in the *G*−*a*−*G* reference sequence, and 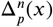 and 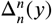, are the neighbor corrections due to preceding (‘p’) and next (ŉ’) residue, respectively. Consequently, the secondary chemical shift for a nucleus *n*, residue *i* is computed from,

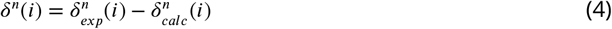

where 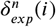 is an experimental resonance assignment belonging to residue *i*.

#### Secondary Chemical Shift for Canonical Secondary Structures

The secondary chemical shifts, expected for fully formed *α*- or *β*-structures, were calculated from,

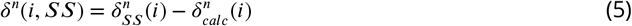

where *δ_SS_* is the expected chemical shift for *α*- or *β*-structures taken from a chemical shift library by Wang and Jardetzky (***Wang and Jardetzky, 2002***); and 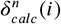 denotes the sequence-specific ‘random-coil’ chemical shift computed from Equation 3.

### NMR Resonance Referencing Offset Corrections

The relative difference between the experimental secondary chemical shifts computed for ^13^*C^α^*, ^13^*C^β^*, and the expected secondary shifts of fully formed *α*- or *β*-structure (***Wang and Jardetzky, 2002***) was used as a measure of a mean referencing error of resonance assignments (***Marsh et al., 2006***). In the current implementation, the effects of fractional deuteration on ^13^*C^α^*, ^13^*C^β^* were not treated explicitly, but assumed to contribute to mean referencing offset. The mean chemical shift referencing offset Δ_*ϵ*_ was computed by minimizing,

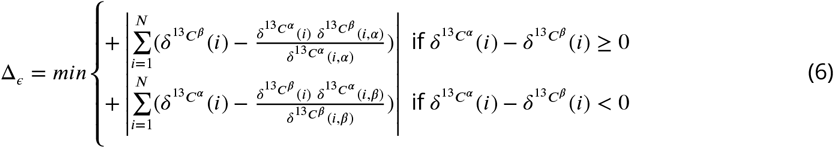

where *δ^n^*(*i, α*) and *δ^n^*(*i, β*) denote the secondary chemical shift in a fully formed *α*- or *β*-structure, respectively.

### Structural Propensity Calculations

The neighbor-corrected Structural Propensity Scores***Tamiola and Mulder (2012)*** Ψ were computed as,

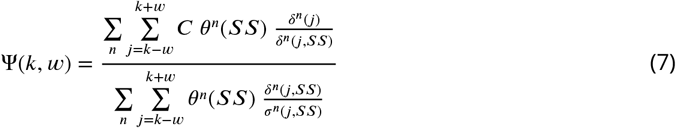

where *δ^n^*(*j*) is the secondary chemical shift of type *n* for the *j*-th residue, *δ^n^*(*j, SS*) represents the expected secondary chemical shift in canonical secondary structure of type *SS*, and *σ^n^*(*j, SS*) is the standard deviation of the expected secondary chemical shift taken from the database by Wang and Jardetzky (***Wang and Jardetzky, 2002***). The parameter *θ^n^*(*SS*) reflects the relative sensitivity of the chemical shift n to secondary structure of type SS. Normalized values of *θ^n^*(*SS*) are given in Table 2.

**Table 2.** Normalized weight parameters *θ^n^*(*SS*), reflecting relative sensitivity of chemical shifts to the canonical secondary structures. *θ^n^*(*SS*) are given in arbitrary units.

| Nucleus | α-helix | β-sheet |
| --- | --- | --- |
| $^1H^N$ | 0.15 | 0.30 |
| $^1H^\alpha$ | 1.00 | 1.00 |
| $^{13}C^O$ | 0.5 | 0.25 |
| $^{13}C^\alpha$ | 1.00 | 1.00 |
| $^{13}C^\beta$ | 1.00 | 1.00 |
| $^{15}N$ | 0.125 | 0.250 |

Secondary structure type discrimination in Equation 7 is achieved by an inclusion of constant C, which is given by Equation 8.

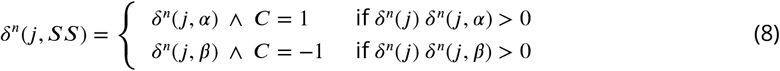

### Sequence Conservation Analysis

Protein sequence data obtained from NMR resonance assignment records were fed to MUSCLE sequence alignment program (***Edgar, 2004***). A column-ordered mean sequence conservation score was used as a measure of sequence similarity across dSPP entries (***Valdar, 2002***). The sequence conservation score was computed assuming BLOSUM62 matrix (***Eddy, 2004***).

### Statistical Analysis

The statistical analysis of protein resonance assignment data was performed in R (version 3.3.2) (***R Core Team, 2013***). In order to avoid data over-binning, the distribution analyses in Figures 1, 2 and 3 were done with variable, sample-dependent bin width. The exact parameters of the histogram analysis are given in Table 3. The residue-specific distributions of structural propensities, depicted on Figure 3b, were computed from the individual kernel density plots of structural propensities with the fixed kernel size of 0.04. The non-parametric single-way ANOVA analysis of theoretical disorder probability distributions was performed using Kruskal-Wallis Test available in R software.

**Table 3.** Parameters for the histogram analysis and plotting of NMR resonance assignment data.

| Property | Samples | Bin width [Unit] |
| --- | --- | --- |
| Temperature | 4509 | 5 [K] |
| pH | 4509 | 0.5 [A.U.] |
| Ionic Strength | 4509 | 50 [mM] |
| Referencing Offset | 4509 | 0.2 [ppm] |
| Sequence Length | 7094 | 25 [A.U.] |
| Structural Propensity | 770653 | 0.04 [A.U.] |

### Secondary structure fraction

The ensemble average secondary structure (SS) fraction was computed from 1000 and 946 PDB files available in *α*S (PED9AAC) (***Fusco et al., 2016***) and MOAG-4 ensembles (kindly provided by Dr. Predrag Kukic, University of Cambridge, UK), respectively. We have used dihedral angle analysis tools of GROMACS (***Berendsen et al., 1995***) to extract residue-specific Φ and Ψ angles. Subsequently, we have computed how many times residues in each ensemble member visited canonical SSs, *α*-helix and *β*-sheet, defined by Φ, Ψ angles of < −48°, −34° > and < −140°, 130° >, respectively. The final fractional SS was computed from arithmetic averages of collected dihedral angle statistics for both ensembles.

### Software Implementation and Availability

The dSPP database was implemented using reactive MeteorJS web application framework with Numerical Python and TensorFlow wrappers. The database is available at https://peptone.io/dspp, both as an interactive application with contextual search and standalone download in JSON format.

## Acknowledgments

Authors acknowledge Dr. Wenwei Zheng (NIDDK, US), Dr. Ruud Scheek (University of Groningen, NL) and Dr. Xavier Periole (Aarhus University, DK) for insightful comments and editorial suggestions. Authors thank Alison Lowndes, Carlo Ruiz and Dr. Adam Grzywaczewski, (NVIDIA Corporation) for facilitating collaboration and access to DGX-1 supercomputer. Jon Wedell (BMRB) is greatly acknowledged for technical support with NMR resonance assignment retrieval from BMRB. We thank Dr. Frans A.A. Mulder (Aarhus University, DK) and Dr. Predrag Kukic (University of Cambridge, UK) for providing structural ensemble models of MOAG-4. Lastly, we want to greatly acknowledge Mark Berger (NVIDIA Corporation) for overwhelming support throughout the execution of this project.

